## Supplementary Material for "Molecular landscape and functional characterization of centrosome amplification in ovarian cancer"

### Supplementary Figures

#### *Supplementary Figure 1 – Spatial heatmaps of centrosome amplification*

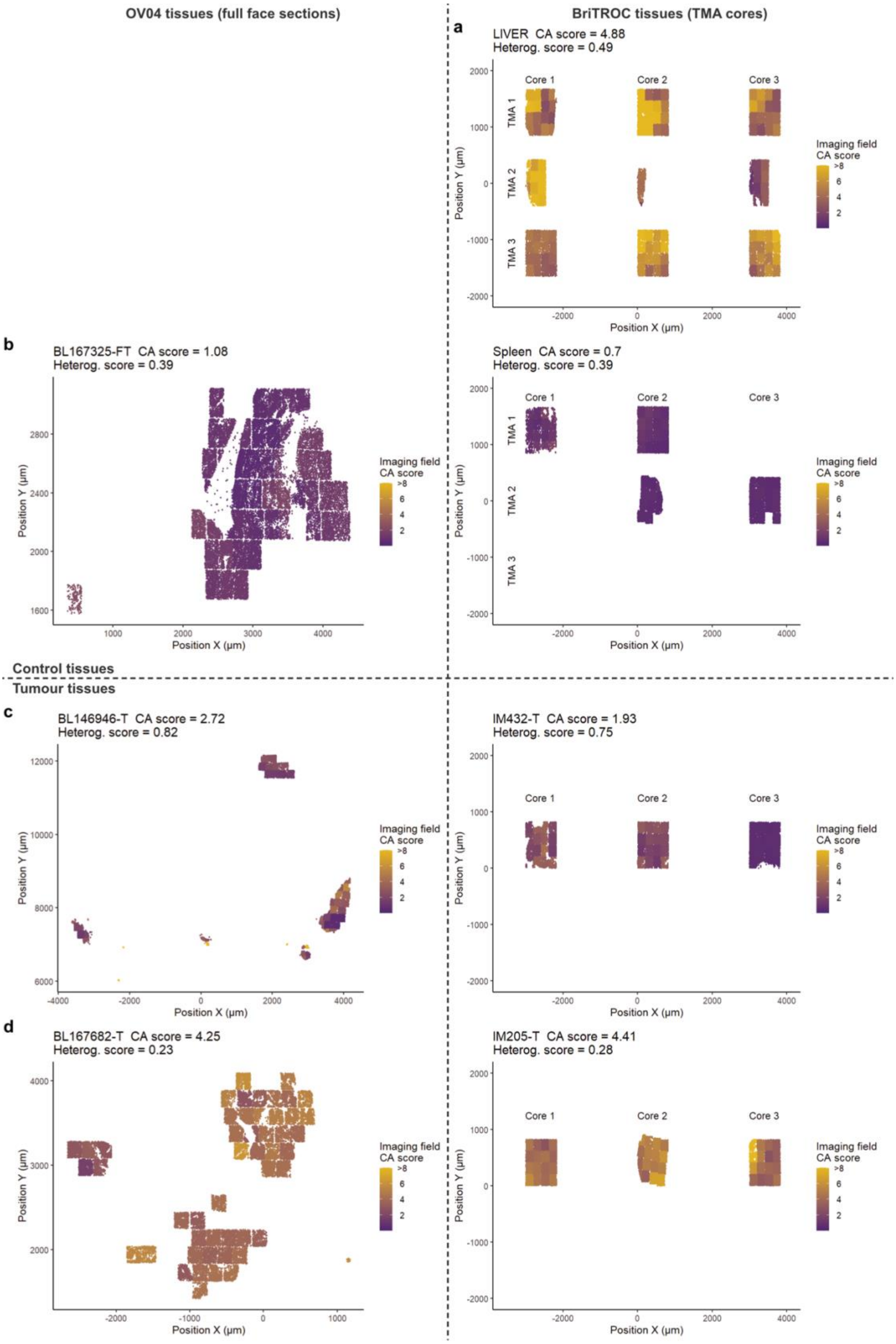

Examples from the OV04 cohort are shown on the left panel, examples from the BriTROC cohort are shown on the right panel. (**a**) Positive control tissue (Liver). (**b**) Negative control (normal) tissues – Fallopian tube on the left, spleen on the right. (**c**) HGSOC tissues with low to mid-range CA and high tissue heterogeneity. (**d**) HGSOC tissues with high CA and low tissue heterogeneity. CA scores are indicated by a colour gradient (purple = low, yellow = high) for each imaging field. Points represent individual nuclei detected during image analyses and were plotted in relation to their physical position (µm) on the microscope slide (x and y axis). *Note that CMYK printing may obscure differences on the CA score colour scale.*

***Supplementary Figure 2 – CA20 gene expression signature is not associated with survival in high CIN cancers.***

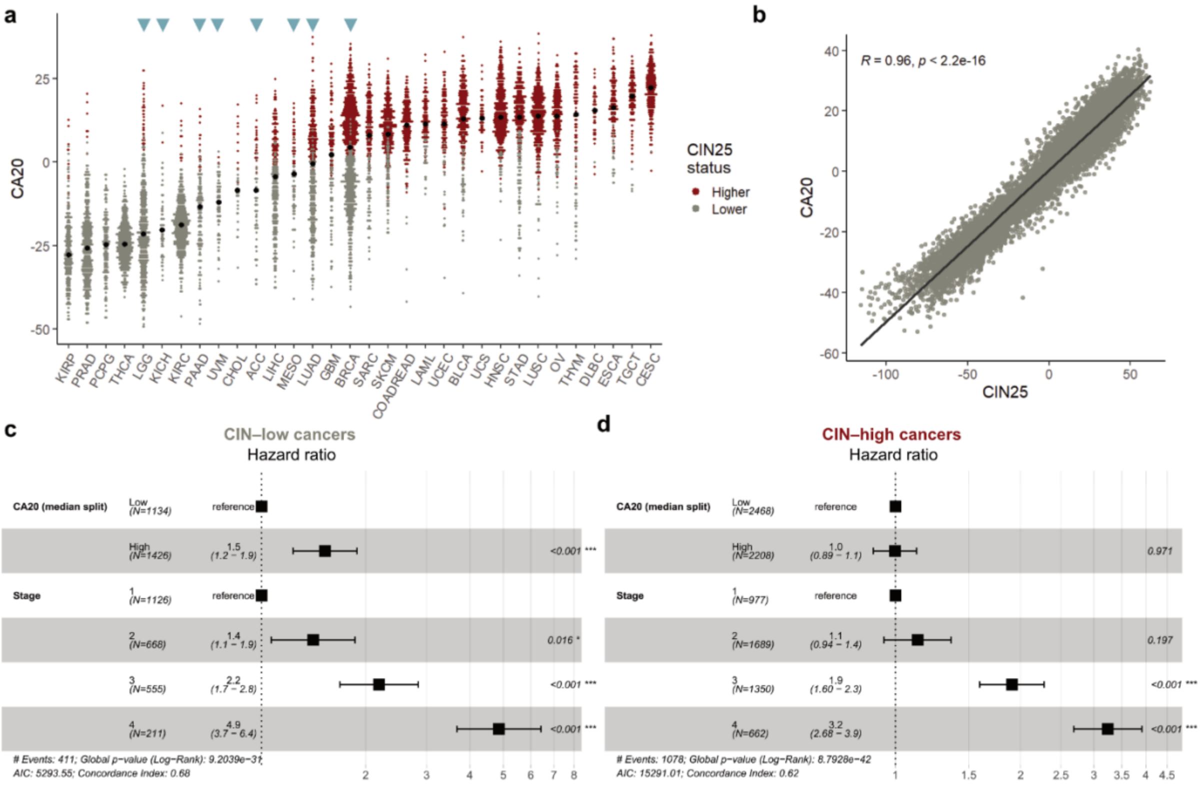

(**a**) CA20 gene expression scores across TCGA pan-cancers (sorted by median CA20 scores indicated by black dots). Each point indicates an individual tumour sample colour coded by the CIN25 gene expression status (median split; red = high, grey = low). Cancer subtypes highlighted by blue arrowheads are cancers in which CA has previously been implicated in poor survival outcome[29]. (**b**) Correlation of CA20 and CIN25 signature expression. (**c**) and (**d**) Forest plots of multivariable Cox proportional hazard modelling on overall survival for low-CIN and high-CIN cancers, respectively.

***Supplementary Figure 3 – Differential gene expression analysis in ovarian cancer vs. normal cell lines.***

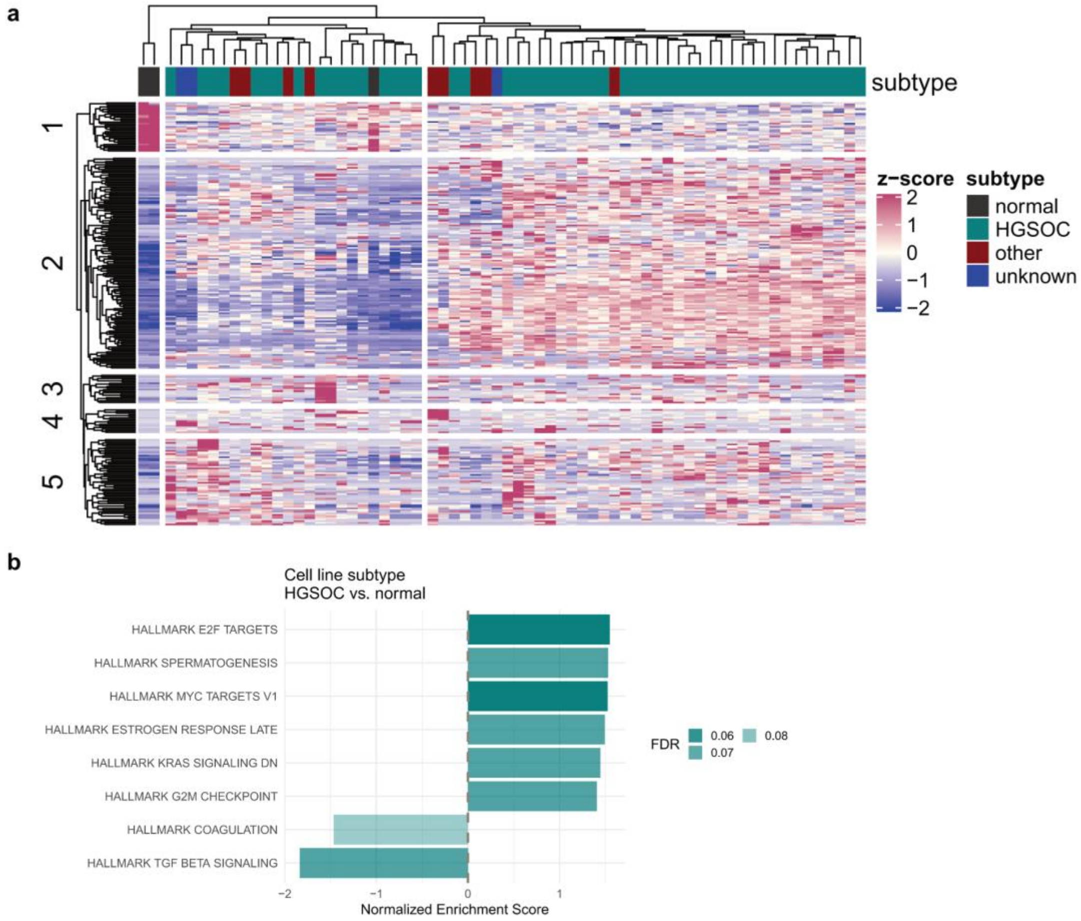

(**a**) Gene expression heatmap across different histological subtypes showing the top 250 differentially expressed genes in HGSOC vs. normal cell lines (FT246, FT194 and IOSE4). Genes and cell lines were grouped using hierarchical clustering. Subtypes are indicated by different colours. (**b**) Gene set enrichment analysis (GSEA) of differentially expressed genes in HGSOC vs. normal cell lines, showing all pathways for which p < 0.05 and FDR < 0.1.

***Supplementary Figure 4 – Centrosome amplification is not associated with distinct genomic features of copy number aberrations in HGSOC tissues.***

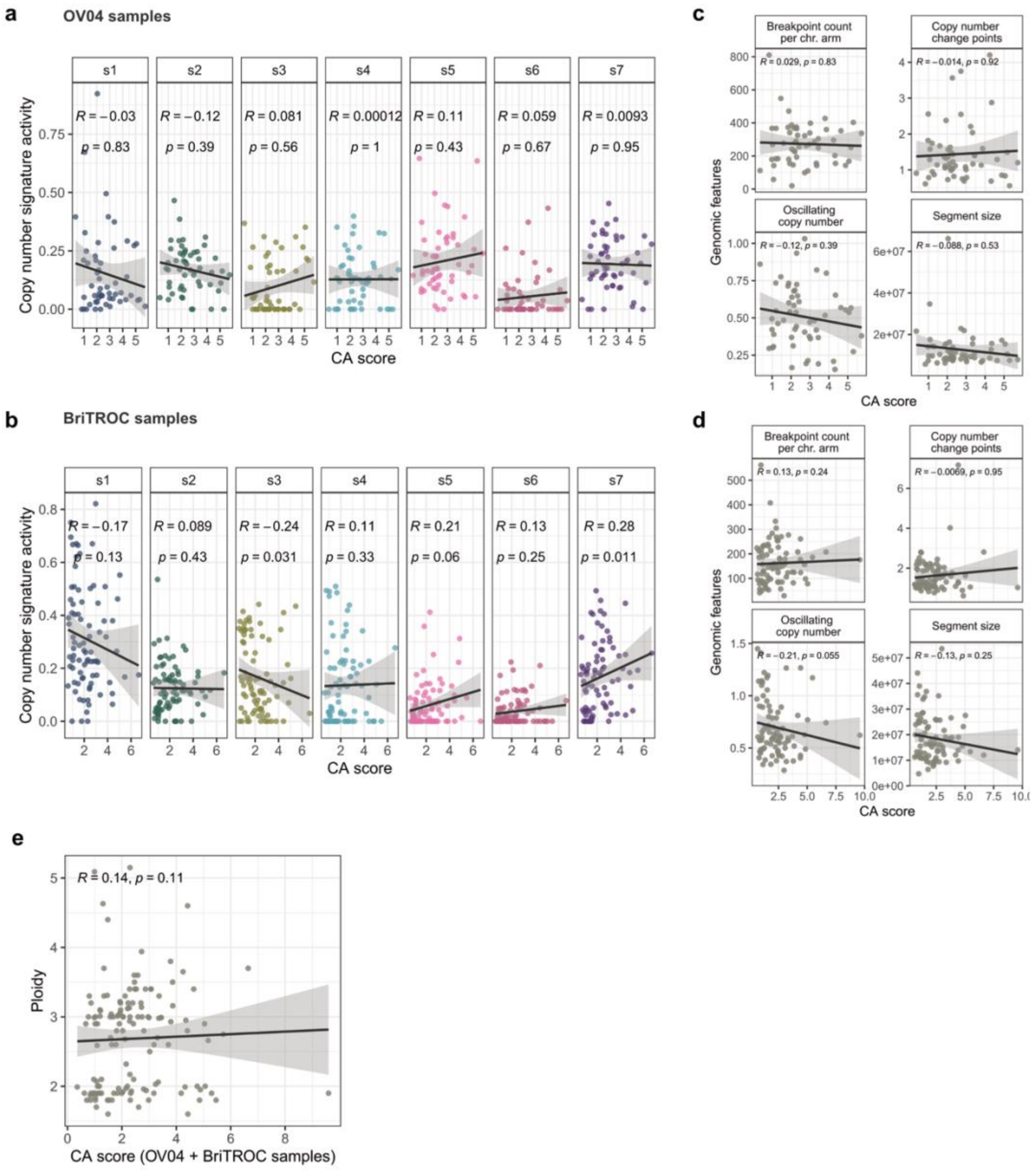

Correlation of copy number signature exposures and CA for (**a**) OV04 samples, and (**b**) BriTROC samples. (**d**) and (**e**) show correlations between CA scores and genomic features for OV04 and BriTROC samples, respectively. (**f**) Correlation of CA scores (from both BriTROC and OV04 samples) and estimated ploidies. Coefficients were estimated using Spearman's rank correlations.

Note that in the BriTROC cohort, there was a negative correlation of CA scores with signature 3, and a positive correlation with signature 7. However, no significant associations were observed following p-value adjustment for multiple comparisons.

***Supplementary Figure 5 – Genomic landscape of ovarian cancer cell lines.***

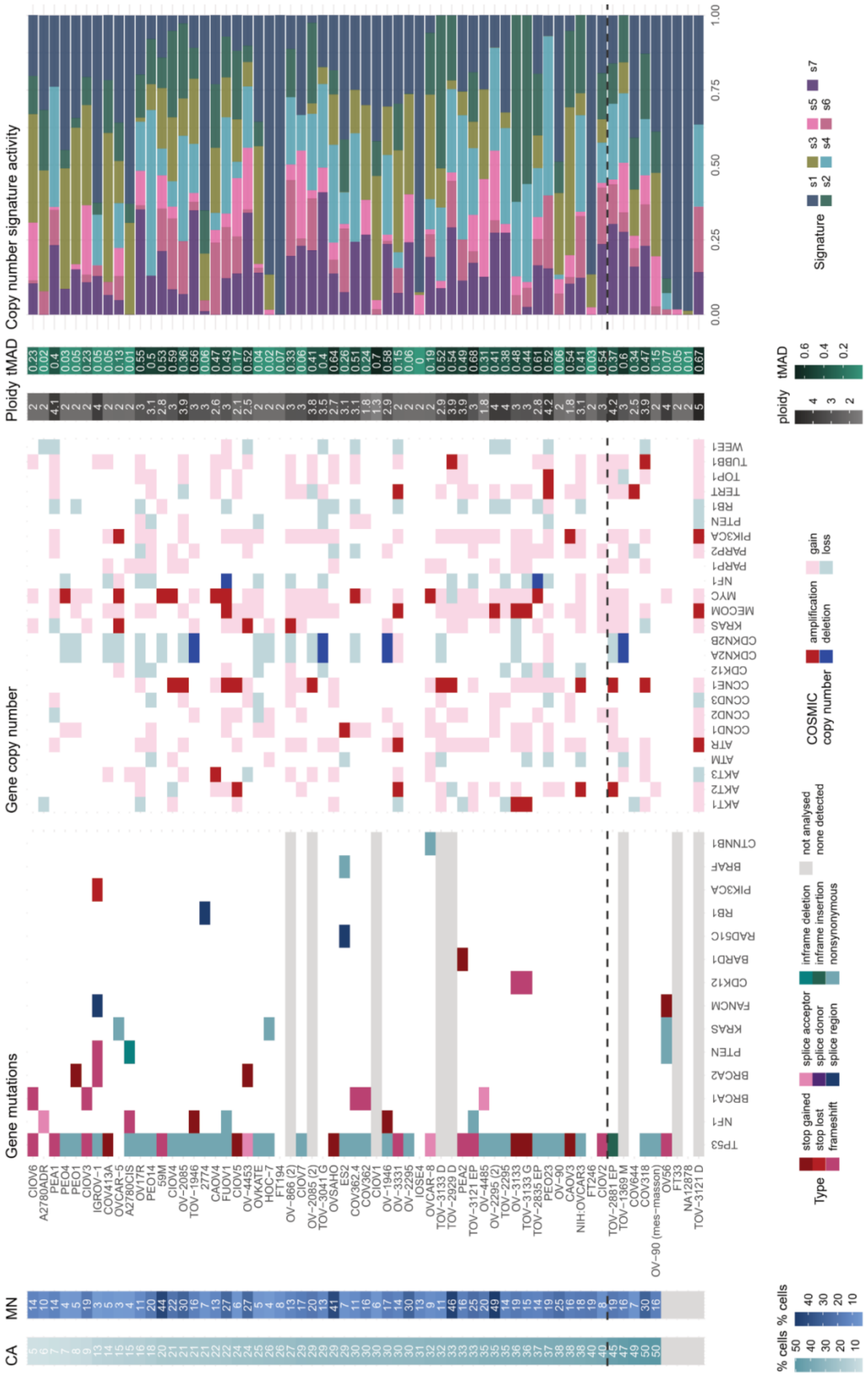

Cell lines are displayed and ordered by their mean centrosome amplification (CA) frequencies (panel 1). Micronuclei (MN) frequencies are plotted along CA frequencies (panel 2). Panel 3 shows an oncoprint plot of gene mutations called from TAm-Seq data. Synonymous and non-pathogenic nonsynonymous mutations or nonsynonymous mutation of unknown clinical significance were excluded. Panel 4 shows gene copy number states (amplification, deletion, gain or loss) for HGSOC genes of interest. Amplifications and deletions were called using the definitions outlined by the Catalogue of COSMIC. Amplifications: ploidy ≤2.7 and CN ≥5; or ploidy >2.7 and CN ≥9. Deletion: ploidy ≤2.7 and CN = 0; or ploidy >2.7 and CN < (ploidy − 2.7). CN gain and losses are defined as a CN change of +1 or −1 from a sample's ploidy, respectively. Ploidies are indicated in panel 5. Trimmed median absolute deviation from CN neutrality (tMAD) scores are shown in panel 6. Panel 7 shows the copy number signature activities derived from ACN-fitted sWGS data for each cell line.

Across all cell lines, the most frequently mutated gene was *TP53* (82%), followed by *NF1* (11%), *BRCA1* (9%) and *BRCA2* (5%). ACN fitting was performed on the sWGS data, and copy number states (amplification, gain, loss and deletion) were called for HGSOC genes of interest depicted in panel 4. The median ploidy across all analysed cell lines was 2.9. The most frequently amplified gene was *CCNE1* (n= 10), followed by *MYC* (n = 9) and *MECOM* (n = 7), while the most frequently gained genes were *MECOM* (n = 30), *PIK3CA* (n = 28) and *MYC* (n = 27). By contrast, copy number deletions were only observed for *CDKN2A*/*B* in four cell lines, and *NF1* in two cell lines. The most frequently lost genes were *RB1* (n = 16), *CDKN2A*/*B* (n = 15) and *NF1* (n = 12). Together, these results suggest that the analysed cell lines are broadly representative of the overall HGSOC case population.

***Supplementary Figure 6 – Correlation of CA and MN with genomic features in ovarian cancer cell lines.***

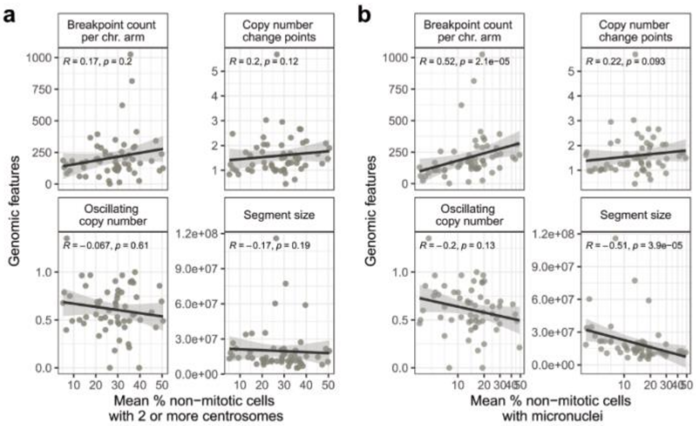

Correlations of genomic features with (**a**) centrosome amplification (CA) and (**b**) micronuclei (MN) frequencies. Coefficients were estimated using Spearman's rank correlations.

### Supplementary Tables

***Supplementary Table 1 – Overview of patients and tissue samples included in centrosome staining study.***

| | | OV04 cohort | BriTROC cohort |
|---|---|---|---|
| Total number of patients | | 85 | 141 |
| Total number of tumour tissues | | 93 | 194 |
| Age at diagnosis | Median | 65 | 65 |
| | Range | 44–83 | 24–85 |
| Histotype | HGSOC | 85 (100%) | 136 (96%) |
| | Endometrioid | | 4 (3%) |
| | Unknown | | 1 (1%) |
| Figo substage | I | 4 (5%) | 10 (7%) |
| | II | 3 (4%) | 15 (11%) |
| | III | 47 (55%) | 90 (64%) |
| | IV | 27 (32%) | 26 (18%) |
| | Unknown | 4 (5%) | |
| Surgery type | Immediate primary surgery (IPS) | 25 (29%) | 92 (65%) |
| | Delayed primary surgery (DPS) | 57 (67%) | 43 (30%) |
| | Unknown | 3 (4%) | 6 (4%) |
| Survival (years) | Median | 2.4 | 4.9 |
| | Range | 0.26–7.7 | 0.76–16.6 |
| Sample type | Full face section | 93 (100%) | |
| | Cores in tissue microarray (TMA) | | 194 (100%) |

***Supplementary Table 2 – Cell line information and culture growth conditions.***

| CellLine | RRID | Media | Supplements | Oxygen | Provider | Type |
|---|---|---|---|---|---|---|
| **2774** | | DMEM/F-12 | 10% FBS | 21% | Jozien Helleman, Netherlands | OV endometrioid carcinoma |
| **59M** | CVCL_2291 | DMEM/F-12 | 10% FBS | 21% | Jozien Helleman, Netherlands | HGSOC |
| **A2780** | CVCL_0134 | RPMI 1640 | 10% FBS + 2mM Sodium Pyruvate | 21% | Jozien Helleman, Netherlands | unknown |
| **A2780ADR** | CVCL_1941 | RPMI 1640 | 10% FBS + 2mM Sodium Pyruvate | 21% | Jozien Helleman, Netherlands | unknown |
| **A2780CIS** | CVCL_1942 | RPMI 1640 | 10% FBS + 2mM Sodium Pyruvate | 21% | Jozien Helleman, Netherlands | unknown |
| **CAOV3** | CVCL_0201 | DMEM/F-12 | 10% FBS | 21% | unknown | HGSOC |
| **CAOV4** | CVCL_0202 | L-15 | 10% FBS | 21% | Jozien Helleman, Netherlands | HGSOC |
| **CIOV1** | | DMEM/F-12 | 10% FBS | 21% | developed in-house; Maria Vias | HGSOC |
| **CIOV2** | | DMEM/F-12 | 10% FBS | 21% | developed in-house; Maria Vias | HGSOC |
| **CIOV3** | | DMEM/F-12 | 10% FBS | 21% | developed in-house; Maria Vias | HGSOC |
| **CIOV4** | | DMEM/F-12 | 10% FBS | 21% | developed in-house; Maria Vias | HGSOC |
| **CIOV5** | | DMEM/F-12 | 10% FBS | 21% | developed in-house; Maria Vias | HGSOC |
| **CIOV6** | | DMEM/F-12 | 10% FBS | 21% | developed in-house; Maria Vias | HGSOC |
| **CIOV7** | | DMEM/F-12 | 10% FBS | 21% | developed in-house; Maria Vias | HGSOC |
| **COV318** | CVCL_2419 | DMEM/F-12 | 10% FBS | 21% | Jozien Helleman, Netherlands | HGSOC |
| **COV362** | CVCL_2420 | DMEM/F-12 | 10% FBS | 21% | Jozien Helleman, Netherlands | OV endometrioid carcinoma |
| **COV362.4** | CVCL_2421 | DMEM/F-12 | 10% FBS | 21% | Jozien Helleman, Netherlands | OV endometrioid carcinoma |
| **COV413A** | CVCL_2422 | DMEM/F-12 | 10% FBS | 21% | Jozien Helleman, Netherlands | OV epithelial-serous carcinoma |
| **COV644** | CVCL_2425 | DMEM/F-12 | 10% FBS | 21% | Jozien Helleman, Netherlands | OV epithelial-mucinous carcinoma |
| **ES2** | CVCL_3509 | McCoy's 5A | 10% FBS | 21% | unknown | HGSOC |
| **FT194** | CVCL_UH58 | DMEM/F-12 | 10% FBS | 21% | Ronnie Drapkin | normal fallopian tube |
| **FT246** | CVCL_UH61 | DMEM/F-12 | 10% FBS | 21% | Ronnie Drapkin | normal fallopian tube |
| **FT33** | CVCL_RK66 | DMEM/F-12 | 10% FBS | 21% | Ronnie Drapkin | normal fallopian tube |
| **FUOV1** | CVCL_2047 | DMEM/F-12 | 10% FBS | 21% | DSMZ, ACC444, Lot7, 7.03.2018 | HGSOC |
| **HeLa** | CVCL_0030 | DMEM/F-12 | 10% FBS | 21% | ATCC | endocervical adenocarcinoma |
| **HOC-7** | CVCL_5455 | RPMI 1640 | 10% FBS + 2mM Sodium Pyruvate | 21% | Jozien Helleman, Netherlands | LGSOC |
| **IGROV1** | CVCL_1304 | RPMI 1640 | 10% FBS + 2mM Sodium Pyruvate | 21% | Jozien Helleman, Netherlands | OV endometrioid carcinoma |
| **IOSE4** | CVCL_0T70 | DMEM/F-12 | 10% FBS | 21% | unknown | normal OV surface epithelium |
| **JHOS-2** | CVCL_4647 | DMEM/F-12 | 10% FBS | 21% | Wendy Fantl, Stanford | HGSOC |
| **Kuramochi** | CVCL_1345 | RPMI 1640 | 10% FBS + 2mM Sodium Pyruvate | 21% | Wendy Fantl, Stanford | HGSOC |
| **NA12878** | CVCL_7526 | RPMI 1640 | 10% FBS + 2mM Sodium Pyruvate | 21% | Caldas lab | lymphoblastoid |
| **NIH:OVCAR3** | CVCL_0465 | RPMI 1640 | 10% FBS + 2mM Sodium Pyruvate | 21% | ATCC | HGSOC |
| **OAW28** | CVCL_1614 | DMEM/F-12 | 10% FBS | 21% | Jozien Helleman, Netherlands | HGSOC |
| **OAW42** | CVCL_1615 | DMEM/F-12 | 10% FBS | 21% | Jozien Helleman, Netherlands | OV cystadenocarcinoma |

***Supplementary Table 2 continued.***

| CellLine | RRID | Media | Supplements | Oxygen | Provider | Type |
|---|---|---|---|---|---|---|
| **OV-1369 (2)** | CVCL_9T12 | OSE | 10% FBS | 5% | Mes Masson, Canada | HGSOC |
| **OV17R** | CVCL_2672 | DMEM/F-12 | 10% FBS | 21% | Jozien Helleman, Netherlands | OV adenocarcinoma |
| **OV-1946** | CVCL_4375 | OSE | 10% FBS | 5% | Mes Masson, Canada | HGSOC |
| **OV-2085** | CVCL_A1SI | OSE | 10% FBS | 5% | Mes Masson, Canada | HGSOC |
| **OV-2085 (2)** | | OSE | 10% FBS | 5% | Mes Masson, Canada | HGSOC |
| **OV-2295** | CVCL_9T13 | OSE | 10% FBS | 5% | Mes Masson, Canada | HGSOC |
| **OV-2295 (2)** | CVCL_9T14 | OSE | 10% FBS | 5% | Mes Masson, Canada | HGSOC |
| **OV-2978** | CVCL_A1SM | OSE | 10% FBS | 5% | Mes Masson, Canada | HGSOC |
| **OV-3133** | CVCL_9T15 | OSE | 10% FBS | 5% | Mes Masson, Canada | HGSOC |
| **OV-3133(2)** | CVCL_9T16 | OSE | 10% FBS | 5% | Mes Masson, Canada | HGSOC |
| **OV-3331** | CVCL_A1SQ | OSE | 10% FBS | 5% | Mes Masson, Canada | HGSOC |
| **OV-4453** | CVCL_9T20 | OSE | 10% FBS | 5% | Mes Masson, Canada | HGSOC |
| **OV-4485** | CVCL_9T21 | OSE | 10% FBS | 5% | Mes Masson, Canada | HGSOC |
| **OV56** | CVCL_2673 | DMEM/F-12 | 10% FBS | 21% | Jozien Helleman, Netherlands | HGSOC |
| **OV-866 (2)** | CVCL_9T22 | OSE | 10% FBS | 5% | Mes Masson, Canada | HGSOC |
| **OV-90** | CVCL_3768 | OSE | 10% FBS | 21% | unknown | HGSOC |
| **OV-90 (mes-masson)** | CVCL_3768 | OSE | 10% FBS | 5% | Mes Masson, Canada | HGSOC |
| **OVCAR-5** | CVCL_1628 | DMEM/F-12 | 10% FBS | 21% | canadian ovarian tissue bank | unknown |
| **OVCAR-8** | CVCL_1629 | RPMI 1640 | 10% FBS + 2mM Sodium Pyruvate | 21% | canadian ovarian tissue bank | HGSOC |
| **OVKATE** | CVCL_3110 | RPMI 1640 | 10% FBS + 2mM Sodium Pyruvate | 21% | unknown | HGSOC |
| **OVSAHO** | CVCL_3114 | RPMI 1640 | 10% FBS + 2mM Sodium Pyruvate | 21% | Simon Langdon | HGSOC |
| **PEA1** | CVCL_2682 | RPMI 1640 | 10% FBS + 2mM Sodium Pyruvate | 21% | Simon Langdon | HGSOC |
| **PEA2** | CVCL_2683 | RPMI 1640 | 10% FBS + 2mM Sodium Pyruvate | 21% | Simon Langdon | HGSOC |
| **PEO1** | CVCL_2686 | RPMI 1640 | 10% FBS + 2mM Sodium Pyruvate | 21% | Simon Langdon | HGSOC |
| **PEO14** | CVCL_2687 | RPMI 1640 | 10% FBS + 2mM Sodium Pyruvate | 21% | Simon Langdon | HGSOC |
| **PEO16** | CVCL_2688 | RPMI 1640 | 10% FBS + 2mM Sodium Pyruvate | 21% | Simon Langdon | HGSOC |
| **PEO23** | CVCL_2689 | RPMI 1640 | 10% FBS + 2mM Sodium Pyruvate | 21% | Simon Langdon | HGSOC |
| **PEO4** | CVCL_2690 | RPMI 1640 | 10% FBS + 2mM Sodium Pyruvate | 21% | Simon Langdon | HGSOC |
| **PEO6** | CVCL_2691 | RPMI 1640 | 10% FBS + 2mM Sodium Pyruvate | 21% | Simon Langdon | HGSOC |
| **SKOV6** | CVCL_A457 | RPMI 1640 | 10% FBS + 2mM Sodium Pyruvate | 21% | Jozien Helleman, Netherlands | cervical squamous cell carcinoma |
| **SNU-119** | CVCL_5014 | RPMI 1640 | 10% FBS + 2mM Sodium Pyruvate | 21% | Wendy Fantl, Stanford | HGSOC |
| **TOV-1369 M** | | OSE | 10% FBS | 5% | Mes Masson, Canada | HGSOC |
| **TOV-1946** | CVCL_4062 | OSE | 10% FBS | 5% | Mes Masson, Canada | HGSOC |
| **TOV-2223 G** | CVCL_4063 | OSE | 10% FBS | 5% | Mes Masson, Canada | HGSOC |

***Supplementary Table 2 continued.***

| CellLine | RRID | Media | Supplements | Oxygen | Provider | Type |
|---|---|---|---|---|---|---|
| **TOV-2295** | CVCL_9T18 | OSE | 10% FBS | 5% | Mes Masson, Canada | HGSOC |
| **TOV-2835 EP** | CVCL_A1SJ | OSE | 10% FBS | 5% | Mes Masson, Canada | HGSOC |
| **TOV-2881 EP** | CVCL_A1SK | OSE | 10% FBS | 5% | Mes Masson, Canada | HGSOC |
| **TOV-2929 D** | CVCL_A1SL | OSE | 10% FBS | 5% | Mes Masson, Canada | HGSOC |
| **TOV-2978 G** | CVCL_9U73 | OSE | 10% FBS | 5% | Mes Masson, Canada | HGSOC |
| **TOV-3041 G** | CVCL_9T24 | OSE | 10% FBS | 5% | Mes Masson, Canada | HGSOC |
| **TOV-3121 D** | | OSE | 10% FBS | 5% | Mes Masson, Canada | HGSOC |
| **TOV-3121 EP** | CVCL_A1SN | OSE | 10% FBS | 5% | Mes Masson, Canada | HGSOC |
| **TOV-3133 D** | CVCL_9T19 | OSE | 10% FBS | 5% | Mes Masson, Canada | HGSOC |
| **TOV-3133 G** | CVCL_4064 | OSE | 10% FBS | 5% | Mes Masson, Canada | HGSOC |
| **TOV-3291 G** | CVCL_9T25 | OSE | 10% FBS | 5% | Mes Masson, Canada | HGSOC |
| **TYK-nu** | CVCL_1776 | DMEM | 10% FBS | 21% | Wendy Fantl, Stanford | HGSOC |
| **TYK-nu.CP-r** | CVCL_3221 | DMEM | 10% FBS | 21% | Wendy Fantl, Stanford | HGSOC |

*Supplementary Table 3 – Primary antibodies for immunofluorescent staining of cell lines.*

| Antibody | Species | Conjugated | Stain1 | Stain2 | Stain3 | Stain4 | Channel | Order number |
|---|---|---|---|---|---|---|---|---|
| Pericentrin | mouse | no | 1 μg/ml | | 1 μg/ml | 1 μg/ml | 647 | ab28144 |
| Pericentrin | rabbit | no | | 1 μg/ml | | | 555 | ab4448 |
| Cep164 | rabbit | no | | | 1 μg/ml | | 555 | 2227-1-AP |
| Centrin 3 | mouse | no | | 1.1 μg/ml | | | 647 | H00001070-M01 |
| pHH3 | rabbit | no | 1 μg/ml | | | | 555 | 06-570 |
| Ck7 - 488 | rabbit | yes | 1 μg/ml | | 1 μg/ml | | 488 | ab208273 |
| Pan-Ck - 488 | mouse | yes | | 1 μg/ml | | | 488 | 53-9003-82 |
| yH2AX - 488 | mouse | yes | | | | 5 μg/ml | 488 | 05-636-AF488 |
| Hoechst | | | 1 μg/ml | 1 μg/ml | 1 μg/ml | 1 μg/ml | 405 | 33342 |

### Supplementary Methods – Method Development

#### *Optimisation of immunofluorescent centrosome staining in clinical FFPE tissues*

FFPE tissues are the most common diagnostic samples available in the clinic. Profiling of centrosome abnormalities in clinical FFPE tissues, however, has previously been hindered by the limited thickness of FFPE sections which prevents imaging of whole cells, and the time-intensive and cumbersome image analysis. The present section describes the development of a reliable high-throughput immunofluorescent microscopy-based assay to unambiguously identify and quantify centrosomes in FFPE samples. This included the optimisation of primary antibody concentrations, detergent concentrations in blocking and antibody dilution buffers, thickness of tissue sections, and antigen retrieval methods. An overview of the immunofluorescent staining workflow for FFPE tissues with optimised conditions (highlighted in blue) for each step is shown in **Methods Fig. 1.**

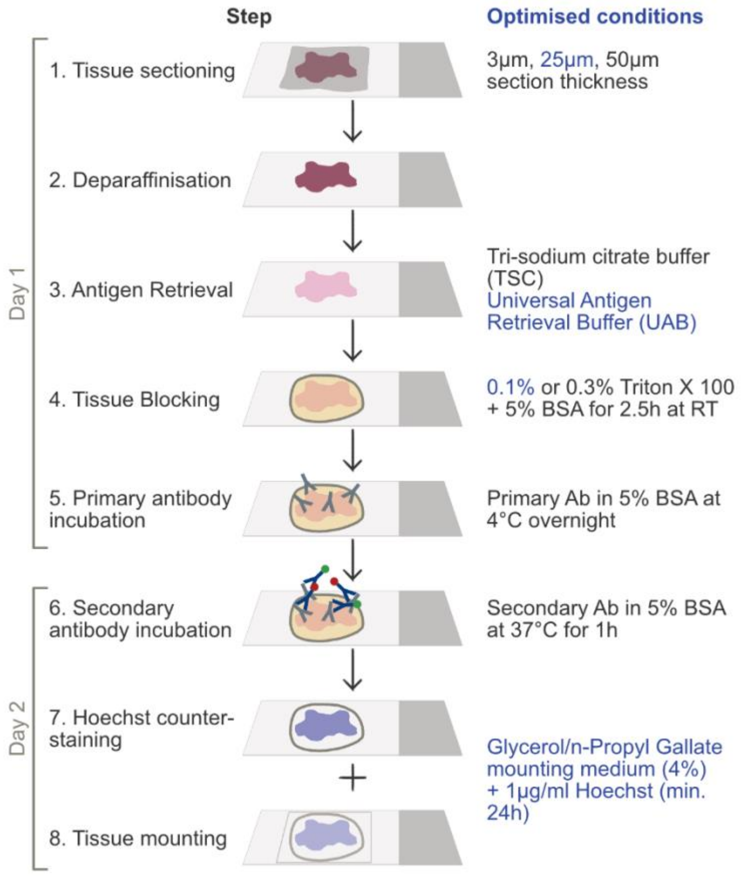

**Methods Fig. 1: Overview of the immunofluorescent staining workflow for FFPE tissue samples** from tissue sectioning through to tissue mounting. Steps not shown here are the crucial washing steps before/after antibody (Ab) incubation and Hoechst staining/tissue mounting. Optimised conditions are highlighted in blue.

1. Optimisation of tissue sectioning thickness

To investigate the effect of tissue section thickness on centrosome detectability, tissue sections of 3µm, 25µm and 50µm thickness were stained using different antigen retrieval methods (see below). As expected, 3µm sections failed to capture intact cells resulting in centrosomes only being detected in a small fraction of cells. By contrast, sections of 50µm thickness failed to provide homogeneous staining results, owing to likely tissue penetration limitations. Using 25µm sections obviated both of these limitations and provided homogeneous staining of mostly intact cells, facilitating the detection of centrosomes in the majority of cells.

2. Optimisation of antigen retrieval methods

Antigen retrieval was performed via heat-induced epitope retrieval (HIER). Two different antigen retrieval buffers were tested: Tri-Sodium Citrate (TSC) and the commercially available Universal Antigen Retrieval Buffer (UAB; ab208572). Both buffers were brought to boil using a conventional microwave. Once the boiling temperature was reached, tissue slides were immersed in either TSC or UAB buffers, boiled for 10 mins at medium-high power and subsequently incubated at room temperature for 25-30 mins. UAB-based antigen retrieval was superior to TSC-based antigen retrieval, and allowed centrosome staining and detection with both, the anti-CDK5RAP2 and anti-Pericentrin antibodies (**Methods Fig. 2**). Importantly, CDK5RAP2 and Pericentrin signal strongly colocalised, while no signal was observed in IgG control stains.

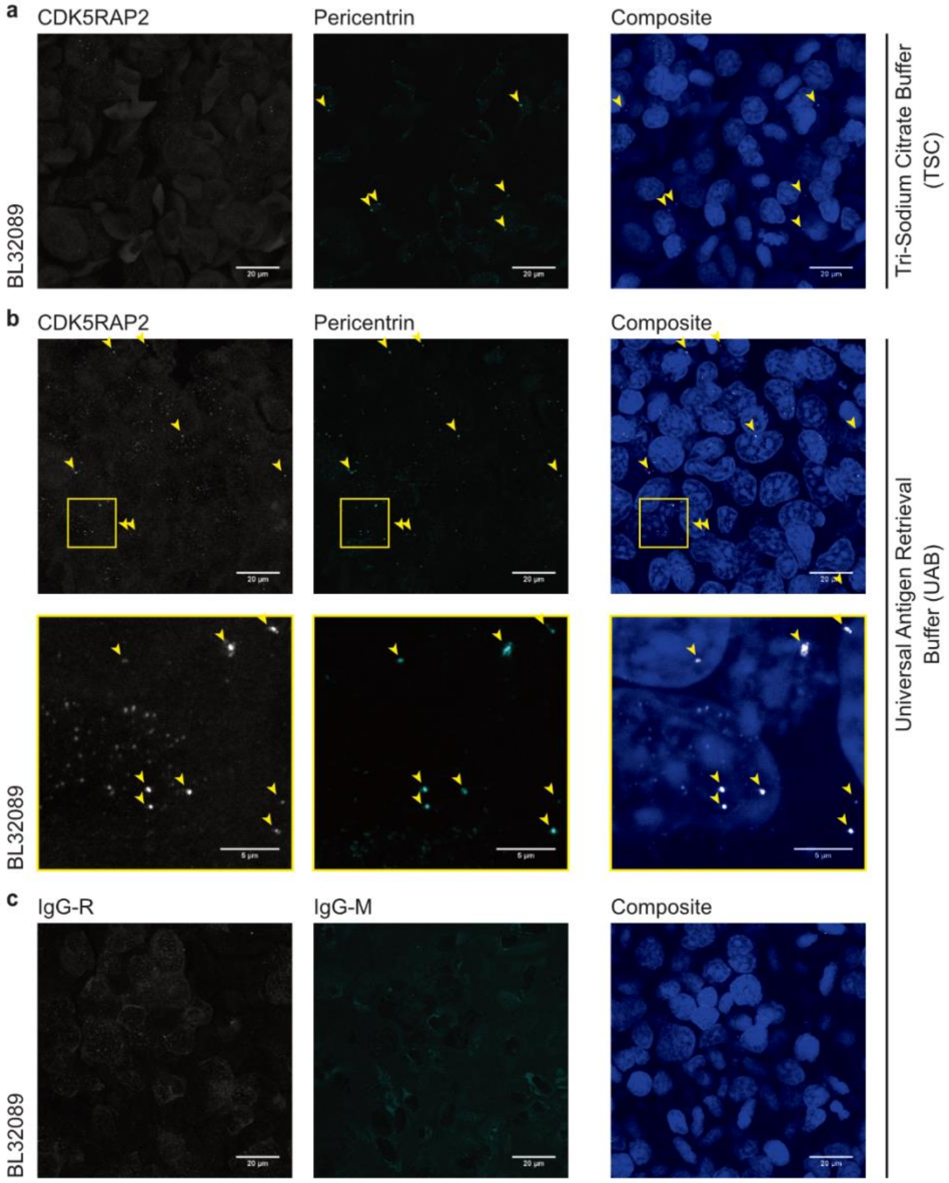

**Methods Fig. 2: The universal antigen retrieval buffer is efficient for centrosome co-staining in thicker FFPE tissue sections**. Tissue sections of 25 µm thickness were co-immunolabelled with anti-CDK5RAP2 and anti-Pericentrin antibodies, and counterstained with Hoechst (blue). Confocal images (max. projections) of centrosome staining using the (**a**) Tri-Sodium Citrate (TSC) or (**b**) Universal antigen retrieval buffer (UAB). (**c**) IgG isotype control (R - rabbit; M - mouse) staining. Scale bar = 20 µm. Zoomed-in scale bar - 5 µm. Centrosomes are indicated by yellow arrow heads.

3. <u>Choice of tissue mounting medium</u>

To preserve stained tissue samples from drying out or deteriorating, samples were mounted in mounting medium and sealed with coverslips following immunofluorescent staining. The main factors considered for the choice of mounting medium included the refractory index of the mountant, fluorophore compatibility, specimen protection and preservation, and the absence of autofluorescent signal originating from the mounting medium. Glycerol/n-Propyl Gallate was found to allow longer-term (>1 month) storage of stained sections and visualisation of nuclei and centrosomes with minimal autofluorescence, and was therefore chosen as mounting medium for tissue sections in this study.

An example of a confocal microscopy image of a tissue stained using the above described and optimised staining workflow is shown in **Methods Fig. 3.**

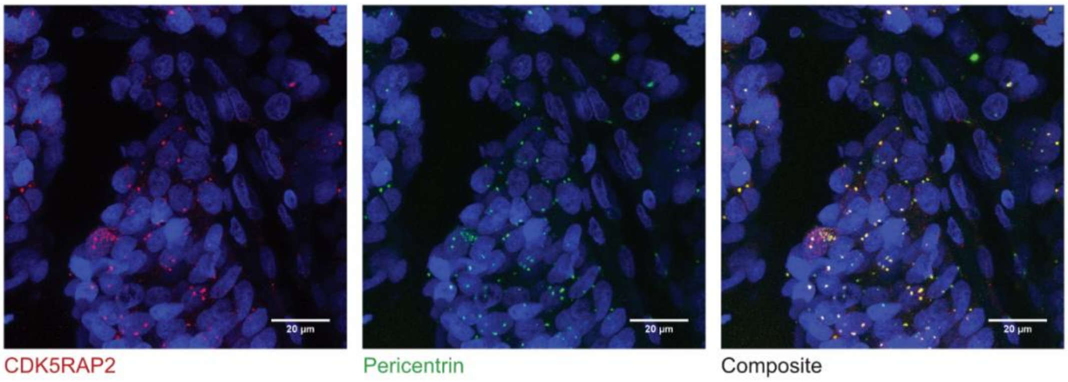

**Methods Fig. 3: Optimised FFPE tissue centrosome staining example**. Confocal image (max. projection) acquired from an FFPE tissue section using the above optimised centrosome staining conditions. Hoechst is shown in blue, CDK5RAP2 is shown in red, Pericentrin is shown in green and colocalising CDK5RAP2 and Pericentrin signal is shown in yellow. Scale bar=20µm.

#### *High-throughput microscopy image acquisition and analysis*

1. <u>Microscopy</u>

In addition to the apparent staining challenges of centrosomes in FFPE tissue samples, three-dimensional imaging and analyses are very time-consuming and cumbersome. Imaging of centrosomes requires high imaging and optical resolution owing to their biologically small size. As a result of these limitations, previous studies investigating centrosome abnormalities in other cancer types have been limited to 3–5 imaging fields capturing only several hundreds of cells per tumour sample.

Using a manual, confocal Leica TCS SP8 microscope at 100× magnification with Z-stacks of 0.5 µm step size and three imaging channels (CDK5RAP2, Pericentrin and Hoechst), image acquisition took approximately 20–30 mins. This means that the collection of imaging data for 10 independent imaging fields would approximately take up to 5 hours for each tissue section (excluding tissue staining and image analysis time), making this approach highly challenging for larger cohort studies.

To circumvent these limitations and to enable higher throughput imaging of centrosomes in HGSOC tissues, we implemented the new Operetta CLS™ high-content analysis system. The Operetta CLS™ spinning disk technology allows high resolution imaging at fast read times and with minimal phototoxicity/photobleaching. In contrast to conventional confocal microscopy, multiple points are scanned simultaneously rather than a single point at a time, facilitating a significantly faster imaging process. An example of three adjacent tissue imaging fields acquired using the Operetta CLS™ system is shown in **Methods Fig. 4**.

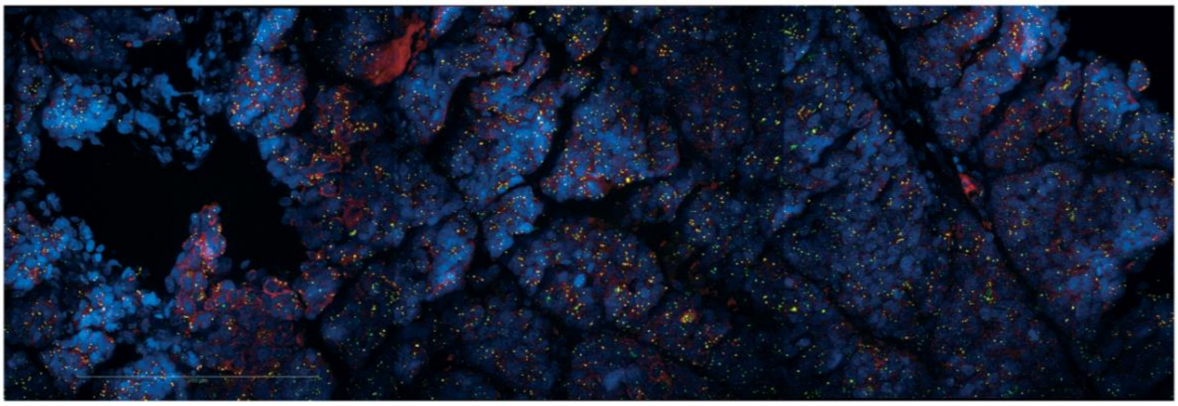

**Methods Fig. 4:** Operetta CLS™ imaging example. Confocal images (max. projections) acquired from an FFPE tissue section using the above optimised centrosome staining conditions. Cells were stained for CDK5RAP2 (red), Pericentrin (green) and DNA (blue). Colocalising CDK5RAP2 and Pericentrin signal is shown in yellow. Scale bar = 200 µm.

To further automate the image acquisition process, we utilised and optimised the intrinsic Operetta PreciScan™ feature (**Methods Fig. 5**).

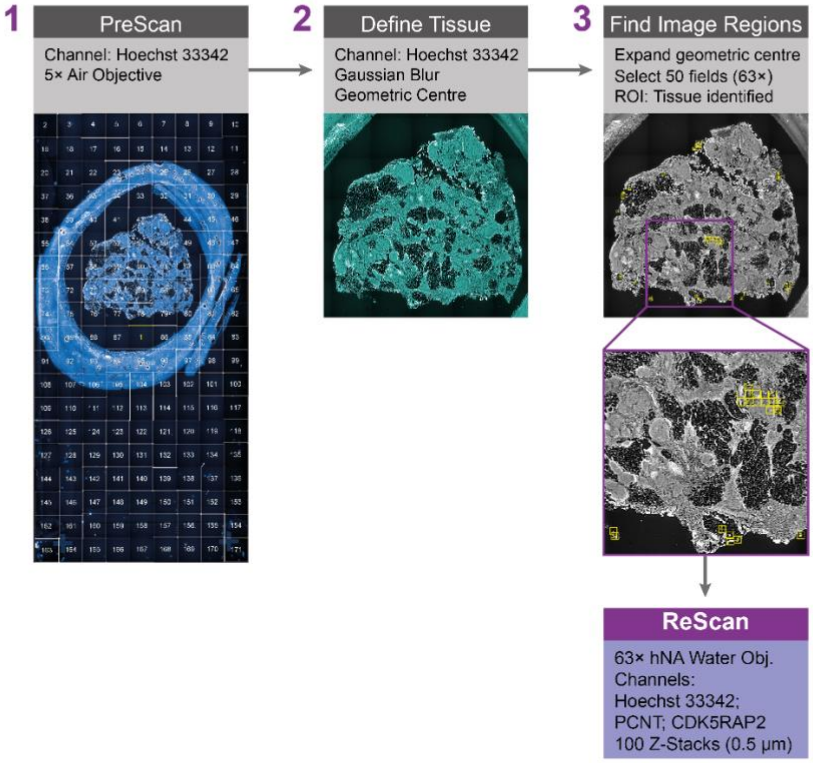

**Methods Fig. 5:** PreScan workflow to identify tissue imaging fields. Flow chart representing the PreScan analysis workflow used to identify tissue imaging fields for subsequent high-resolution ReScanning. Following low magnification (5×) imaging of whole tissue slides, global images were filtered using Gaussian blur and tissue was identified based on Hoechst 33342 signal. The geometric centre of the tissue was then calculated and expanded to create a region of interest covering the majority of the tissue. Following morphological properties calculations, 50 imaging fields were placed within the identified tissue for subsequent high resolution ReScan (63× magnification).

The PreciScan™ plug-in allows a fully automated workflow of a low magnification (5×) pre-scan, followed by an image analysis step to identify tissue regions of interest, and finally higher magnification (63×) re-scans of selected imaging fields. This approach allowed scanning of 50 independent non-overlapping imaging fields for each tissue section within a reasonable time frame, providing a larger dataset and better representation of tumour tissues.

2. <u>Image Analysis</u>

Images and data were reconstructed and analysed using the Harmony High-Content Analysis software. Following nuclei segmentation and spot counting, centrosome amplification (CA) scores were calculated for each of the 50 fields within a tissue sample using the following formula:

$$\text{CA Score} = \frac{\sum Centrosomes\ per\ field}{\sum Nuclei\ per\ field}$$

**Methods Fig. 6** illustrates the workflow of the optimised image analysis sequence used for the quantification of centrosomes in HGSOC FFPE tissue samples.

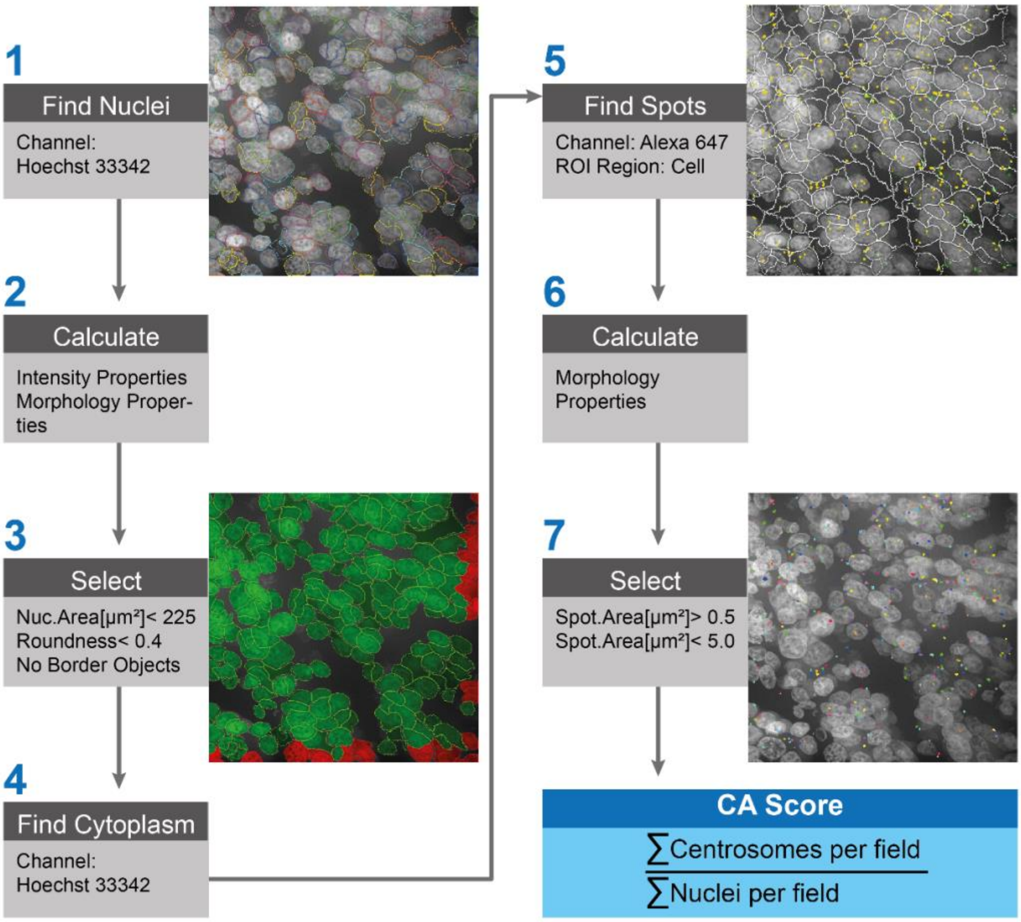

**Methods Fig. 6:** Image Analysis Workflow. Flow chart representing the image analysis workflow used to identify and quantify centrosomes in human FFPE tissues. Following image acquisition, nuclei were segmented and selected based on size and roundness. Border objects were removed to only include whole nuclei during image analysis. Cytoplasm was identified based on diffusing Hoechst signal and used to identify cell boundaries. Cell boundaries were then used as a region of interest (ROI) within which centrosomes were identified and filtered based of morphology properties. Tissue-wide CA scores were estimated as the median CA score across all imaging fields.

3. Quality control for CA analyses

The large amount of data collected (>3.5 × $10^6$ images across all tissue samples) prohibited manual quality control and inspection of all imaging data. To improve the robustness of our approach, we randomly selected ten tissue sections that were stained with secondary antibodies only (no primary antibodies) to estimate background noise, autofluorescence, and the rate of false positive detection of centrosomes (**Methods Fig. 7a**). Based on these experiments, all imaging fields with estimated centrosome amplification scores (CA score) < 0.1 were removed from downstream analyses (**Methods Fig. 7b**). Inspection of these imaging fields showed that they contained images that were out of focus or captured areas of hydrophobic pen (used during antibody staining) that were misidentified as tissue during the PreScan workflow.

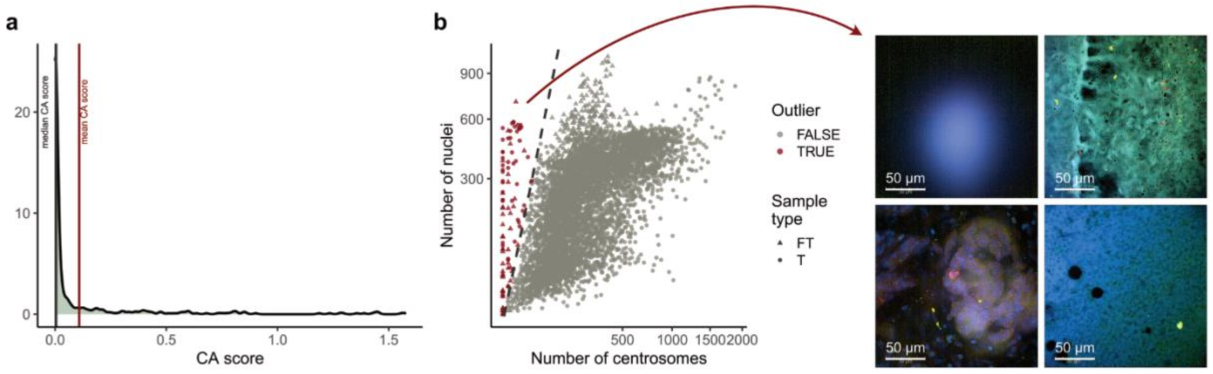

**Methods Fig. 7: Removal of outlier imaging fields.** (**a**) Density plot of CA scores determined from ten control HGSOC tissue sections stained with secondary antibodies only. Median and mean CA scores across all ten sections are indicated by vertical grey and red lines, respectively. (**b**) Scatter plot showing number of nuclei vs number of centrosomes detected in each imaging fields. Low outliers are highlighted in red. Dashed diagonal line indicates CA score = 0.1. Example images from outlier imaging fields are shown on the right. T = Tumour; FT = Fallopian tube. Scale bars = 50 µm.
